# Nutrient profiling reveals extracellular uridine as a fuel for pancreatic cancer through uridine phosphorylase 1

**DOI:** 10.1101/2021.06.07.447448

**Authors:** Matthew H. Ward, Zeribe C. Nwosu, Pawan Poudel, Steven Kasperek, Zach Tolstyka, Rosa E. Menjivar, Chanthirika Ragulan, Gift Nyamundanda, Li Zhang, Anthony Andren, Christopher J. Halbrook, Eileen S. Carpenter, Marina Pasca di Magliano, Anguraj Sadanandam, Costas A. Lyssiotis

## Abstract

Pancreatic ductal adenocarcinoma (PDA) is a lethal disease characterized by high invasiveness, therapeutic resistance, and metabolic aberrations. Although altered metabolism drives PDA growth and survival, the complete spectrum of metabolites used as nutrients by PDA remains largely unknown. Here, we aimed to determine novel nutrients utilized by PDA. We assessed how >175 metabolites impacted metabolic activity in 19 PDA cell lines under nutrient-restricted conditions. This analysis identified uridine as a novel metabolite driver of PDA survival in glucose-deprived conditions. Uridine utilization strongly correlated with expression of the enzyme uridine phosphorylase 1 (UPP1). Metabolomics profiling, notably ^13^C-stable isotope tracing, revealed that uridine-derived ribose is the relevant component supporting redox balance, survival, and proliferation in glucose-deprived PDA cells. We demonstrate that UPP1 catabolizes uridine, shunting its ribose component into central carbon metabolism to support glycolysis, the tricarboxylic acid (TCA) cycle and nucleotide biosynthesis. Compared to non-tumoral tissues, we show that PDA tumors express high *UPP1*, which correlated with poor overall survival in multiple patient cohorts. Further, uridine is enriched in the pancreatic tumor microenvironment, and we demonstrate that this may be provided in part by tumor associated macrophages. Finally, we found that inhibition of *UPP1* restricted the ability of PDA cells to use uridine, and that *UPP1* knockout impairs tumor growth *in vivo*. Our data identifies uridine catabolism as a critical aspect of compensatory metabolism in nutrient-deprived PDA cells, suggesting a novel metabolic axis for PDA therapy.

## Introduction

Despite decades of research and drug development, pancreatic cancer remains one of the two deadliest cancers (Siegel et al., 2019). Pancreatic ductal adenocarcinoma (PDA) represents the vast majority of pancreatic cancer diagnoses (Gordon-Dseagu et al., 2018; Ryan et al., 2014; Wolfgang et al., 2013), and its high mortality is attributed both to a preponderance of late diagnoses and a lack of effective treatments (Biancur and Kimmelman, 2018; Hidalgo, 2010).

PDA biology is driven by microenvironmental and intrinsic variables that impact therapy and tumor aggressiveness. The PDA tumor microenvironment (TME) contributes to therapeutic resistance and is characterized by the secretion of extracellular matrix (ECM) from stromal fibroblasts, an increase in the pressure inside the pancreas, and localized destabilizations of the vascular pressure leading to the collapse of arterioles and capillaries (Chu et al., 2007; Provenzano et al., 2012; Puleo et al., 2018). This phenomena contributes to therapeutic resistance and results in metabolic heterogeneity within the tumor (DuFort et al., 2016). Specifically, areas with low oxygen saturation and abnormal nutrient profiles develop unevenly throughout the tumor and promote metabolic adaptation at the cellular level (Kamphorst et al., 2015; Koong et al., 2000; Sullivan et al., 2019).

PDA cells surviving nutrient and oxygen deficiencies exhibit metabolic adaptations that increase their scavenging and catabolic capabilities. Such adaptations include autophagy, macropinocytosis, and the upregulation of nutrient transporters, which allow the scavenging of critical metabolites from extracellular and intracellular sources, and all support PDA cell growth (Commisso et al., 2013; Kamphorst et al., 2015; Yang et al., 2011; Zhao et al., 2016). Furthermore, recent studies have defined additional nutrient sources, including ECM, immune, and stromal derived metabolites (Halbrook et al., 2019; Kim et al., 2020; Sousa et al., 2016). These mechanisms allow metabolic crosstalk between cell types and thus the symbiotic exchange of nutrients to support tumor metabolic homeostasis and growth under nutrient-deficient conditions.

While these isolated examples of nutrient exchange have been uncovered, comprehensive screens with the power to identify many such mechanisms have not been previously performed. Here, we used Biolog Phenotype Microarrays to measure the ability of a large panel of PDA cell lines to catabolize over 175 different potential energy sources. Our goal was to identify candidate metabolites that support PDA survival. We find that exogenous uridine supports PDA metabolic homeostasis and growth during glucose deprivation, a phenotype that is driven by uridine phosphorylase 1 (UPP1). *UPP1* is highly expressed in PDA tumors relative to non-tumors, and high *UPP1* expression is associated with a worse overall survival. Uridine is available in the pancreatic interstitial fluid, and we demonstrate that this is released by tumor educated macrophages. Mechanistically, *UPP1* enables the catabolism of the ribose moiety from uridine through central carbon metabolism and its recycling for nucleotide salvage. We also show that targeting *UPP1* in a xenograft PDA tumor model selectively impaired tumor growth. These data support that uridine catabolism drives PDA compensatory metabolism and malignancy, thus representing a potential therapeutic target.

## Results

### Nutrient screening in a panel of PDA cells identifies extracellular uridine as a novel fuel

To screen for metabolites that could fuel metabolism in nutrient-deprived PDA cells, we applied the Biolog metabolite phenotypic screen platform (**Fig. 1a**) on a panel of 20 PDA human cell lines (19 after quality control) and two immortalized non-malignant pancreas cell lines (hPSC and HPNE cells). The Biolog assay assessed cellular ability to capture and metabolize >175 nutrients in an array format under nutrient limiting conditions (0mM glucose, 0.3mM glutamine, 5% dialyzed fetal bovine serum). The panel of nutrients includes various carbon energy and nitrogen substrates, such as carbohydrates, nucleic acids, glycosylamines, metabolic intermediates, amino acids, and dipeptides (**Table 1**). Metabolic activity was assessed by monitoring the reduction of a tetrazolium-based dye, a readout of cellular reducing potential, every 15 minutes over a period of 74.5 hours. The kinetic measurements evaluated several parameters, including the time taken for cells to adapt to and catabolize a nutrient (lambda), the rate of uptake and catabolism (mu or slope), the total metabolic activity (AUC or area under the curve), and the maximum metabolic activity (**Fig. S1a-d**). Analyses of global nutrient consumption profiles revealed that PDA cells cluster into 2-4 subsets with varying metabolite utilization capabilities (**Fig. 1b, Fig. S1e**). While there was a variation in the use of metabolites such as α-methyl-D-galactoside, methyl pyruvate, D-galactose, D-malic acid and citric acid between cell lines (**Fig. 1c**), as well as many dipeptides (**Fig. S1f**), there was less heterogeneity in the use of other metabolites such as uridine, glucose, fructose, maltose and glycogen (**Fig. S1g**), indicating metabolites utilized by most of the cell lines. Of these, uridine was intriguing given that rather than being a carbohydrate, it is a nucleoside.

**Figure 1.**
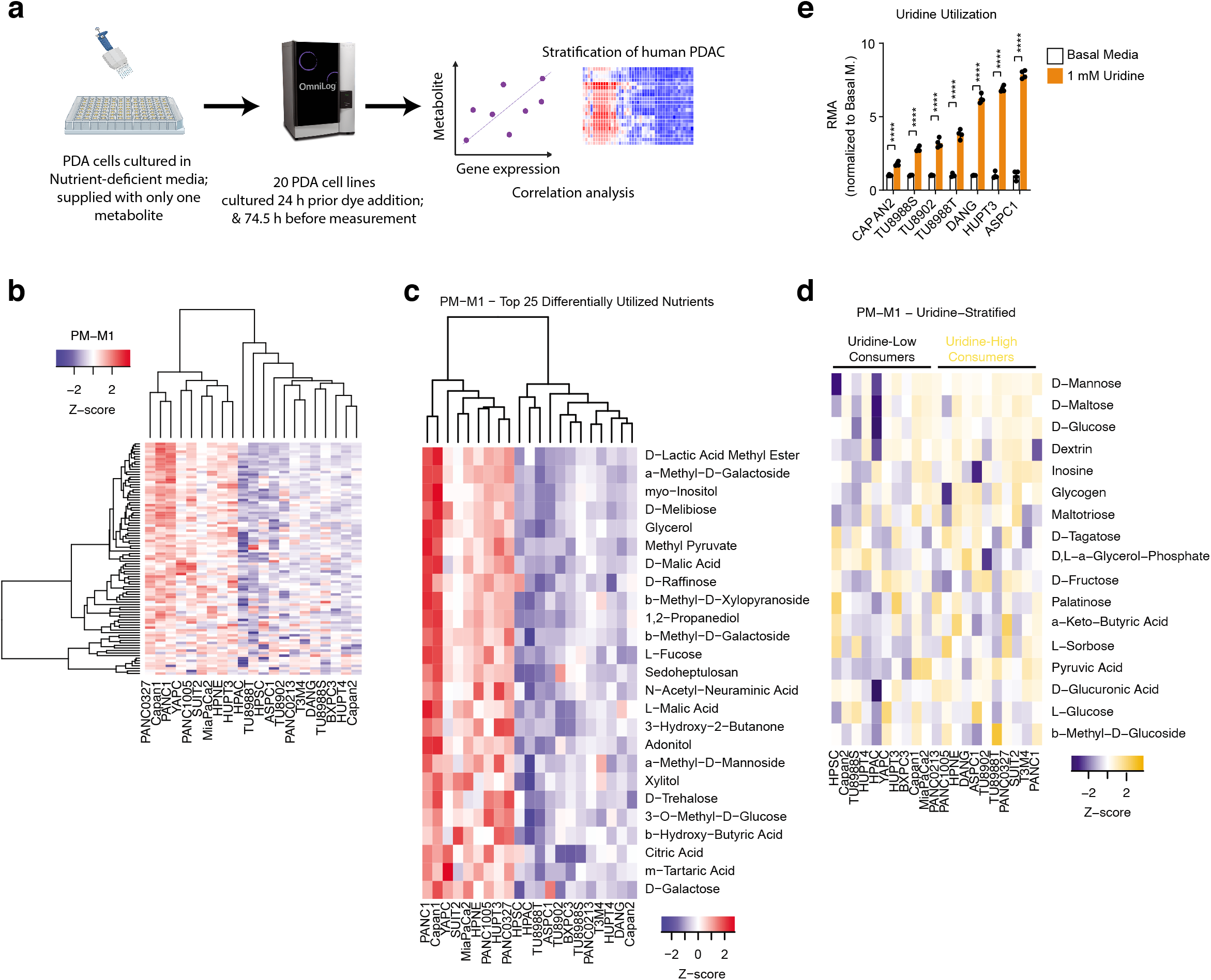
Profiling of PDA metabolite utilization identifies uridine. **a**. Schematic representation of metabolite utilization analysis workflow. Nutrient utilization profile as measured by Biolog (using the Omnilog device) was used to determine metabolite consumption that correlates with gene expression. **b.** Heatmap showing the differential use of metabolites by 19 PDA cells in nutrient-depleted media over 74.5 h culture. Red denotes over-utilization; blue denotes under-utilization. PM-M1 plate contains mainly carbohydrates. **c.** Heatmap showing metabolites that were utilized mainly a subset of cell lines. **d**. Heatmap showing PDA cell lines stratified into the high and low uridine consumers. **e**. Measurement of relative metabolic activity (RMA) of PDA cell lines cultured in media supplemented with 1 mM uridine for 74.5 h. Basal media indicates SILAC RPMI 1640 supplemented with 5% dFBS, 0.3mM Gln, 1.148mM Arg, and 0.2189 mM Lys.

To determine the underlying features of uridine metabolism in PDA cells, we further stratified the cell lines into high and low metabolizers of uridine. We found that cells that consumed a higher level of uridine also consumed considerably more glucose (**Fig. 1d**). Using representative low and high-uridine consuming cells, we confirmed that supplementation with uridine enhanced the metabolic activity of PDA cell in nutrient-deprived media (**Fig. 1e**). These data identify uridine among several other metabolites that could serve as alternative nutrient sources for nutrient-restricted PDA cells.

### UPP1 expression correlates with uridine consumption and is high in PDA tumors

To determine the molecular features underlying uridine consumption, we compared the gene expression profile of uridine-high and low consumers using the Cancer Cell Line Encyclopedia (CCLE) data (**Fig. 2a**). We identified >700 differentially expressed genes (DEGs, *P*<0.05) (**Supplementary Table 1**). Among the topmost downregulated genes in high uridine consumers were insulin-like growth factor 1 receptor (*IGF1R*), trefoil factor 2 (*TFF2*, known to inhibit gastric motility and secretion), and glutathione s-transferase zeta 1 (*GSTZ1*) (**Fig. 2a**). In contrast, the highly expressed genes in uridine-high consumers were *HUS1* (in cell cycle), *PPFIA1*, mitochondrial folate transporter *SLC25A32*, cryptochrome circadian regulator 1 (*CRY1*) and uridine phosphorylase 1 (*UPP1*) (**Fig. 2a**). Of these, *UPP1* is the most directly related to uridine. *UPP1* encodes an enzyme that mediates the catabolism of uridine to uracil and ribose-1-phosphate and has not been previously characterized in PDA.

**Figure 2.**
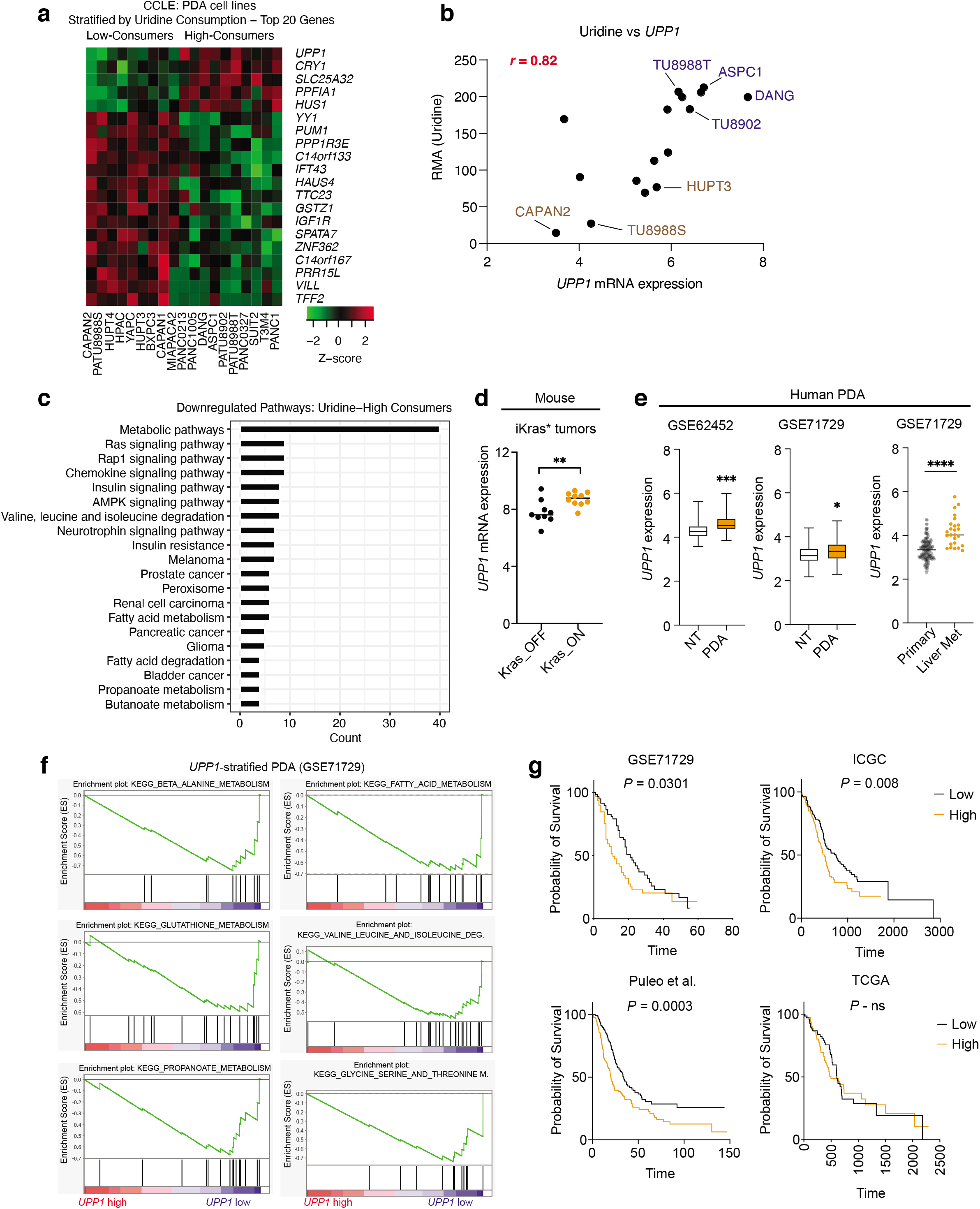
UPP1 expression correlates with uridine metabolism in PDA. **a.** Topmost genes from the cancer cell line encyclopedia PDA cells stratified based on uridine consumption as determined from the metabolite utilization (Biolog) profiling in Fig. 1d. **b**. Spearman correlation between relative metabolic activity (RMA) derived from uridine catabolism and UPP1 transcriptional expression in 16 PDA cell lines with corresponding data in the analyzed dataset.. **c**. Functional annotation analysis showing the pathways downregulated in UPP1-high PDA cell lines. **d**. Expression of *Upp1* in murine inducible oncogenic kras (iKras*) tumors published in the dataset GSE32277. **e**. *UPP1* expression in PDA tumors relative to non-tumoral pancreatic tissue samples in the indicated gene datasets. **f**. Gene set enrichment analysis, showing downregulated pathways in UPP1-high PDA. The genes used for the analysis were derived from the PDA dataset GSE71729 stratified into UPP1-low (n=72) and UPP-high (n=73) subsets. **g**. Kaplan Meier overall survival analysis (log-ranked test) based on *UPP1* expression in the indicated datasets (see Methods). TCGA – the cancer genome atlas; NS – not significant.

Complementary to the CCLE gene correlation data, *UPP1* also emerged among the topmost metabolic genes correlated with their respective metabolites in our comprehensive analysis of the Biolog data (**Supplementary Table 2**) using an independent previously published dataset (Klijn et al., 2015). There was a strong, and near linear, correlation of uridine utilization and *UPP1* expression among the cell lines in our panel (*r* = 0.82, *P*=0.0001, **Fig. 2b**), which was further confirmed by qPCR mRNA expression analysis from cells profiled under nutrient-deficient conditions, as relevant to the Biolog assay (**Fig. S2a**). To determine the specificity of the association between uridine catabolism and *UPP1* expression, we assessed the correlation of *UPP1* expression to other nucleosides included in the Biolog assays. While both inosine and adenosine were avidly catabolized, adenosine, inosine, and thymidine were effectively not correlated with *UPP1* expression (**Fig. S2b**). These results indicate that the association between *UPP1* expression and uridine catabolism is both robust and specific, and that the catabolic mechanism(s) of other nucleosides is independent of UPP1.

To gain insight on the molecular implications of UPP1, we analyzed pathway annotation of DEGs in UPP1-high cells. The upregulated genes in UPP1-high cell lines included those involved in endocytosis (e.g., *RAB10, RAB7A, PARD6B, NEDD4, CBL, EGFR*), and several inflammation/immune-related pathways, notably NFkB signaling (e.g., *CHUK, DDX58, TRIM25, TAB2, RELA*) (**Supplementary Table 3**) – potentially linking UPP1 to TME activities. In contrast, UPP1-high cell lines showed a profound downregulation of metabolic pathways (**Fig. 2c**), a pattern that is consistent with vulnerable metabolic activities in cancer. In addition, in line with metabolic impairment, gene ontology analysis revealed a notable downregulation of genes in the cellular component ‘mitochondria’ in UPP1-high cell lines, as compared to cytosolic and membrane signatures for the upregulated genes (**Fig. S2c**).

To further establish the potential relevance of UPP1 in PDA, we analyzed its expression in microarrays of mouse PDA cell lines and tumor datasets. Notably, using datasets from our inducible Kras mouse model of PDA (Collins et al., 2012; Ying et al., 2012), we evaluated the Kras-mediated regulation of *Upp1*. Mutant Kras expression, the signature driving oncogene in PDA, led to a high UPP1 expression *in vitro* (**Fig. S2d)** and in a subcutaneous xenograft model *in vivo* (**Fig. 2d**). In human PDA, *UPP1* was among our recently identified genes of high priority (Nwosu et al., 2021). UPP1 is highly expressed in PDA tumors compared to non-tumoral samples, as well as in liver metastasis compared to primary tumors (**Fig. 2e**). Further, similar to the UPP1-high cell lines, UPP1-high patient tumors displayed a general downregulation of metabolic pathways (**Fig. 2f**) with the obvious exception of glycolysis (shown in **Fig. 5a**), and also showed a prominent upregulation of inflammatory/immune processes (**Fig. S2e**), and a differential expression of tricarboxylic acid (TCA) cycle genes that are fundamental to mitochondrial function (**Fig. S2f**). In multiple patient cohorts, the high expression of *UPP1* predicted poor overall survival outcome (**Fig. 2g**). Collectively, this evidence establishes, for the first time, that UPP1 is a critical metabolic enzyme in PDA.

#### Uridine-derived ribose supports pancreatic cancer metabolism

We found that exogenous uridine can support the metabolism of PDA cells cultured in nutrient-depleted, glucose-free conditions, in a manner similar to glucose, when examined at equimolar concentrations (**Fig. 3a**). Based on this observation, we hypothesized that UPP1 liberated ribose from uridine, and that this ribose was recycled into glycolysis. To test this hypothesis, we supplemented cells grown in glucose-free media with ribose, a cell-permeable substitute for ribose-1-phosphate, to determine if this was sufficient to replicate the effects of exogenous uridine. Indeed, supplementation with ribose alone fueled the generation of reducing potential in the form of NADH (**Fig. 3a**).

**Figure 3.**
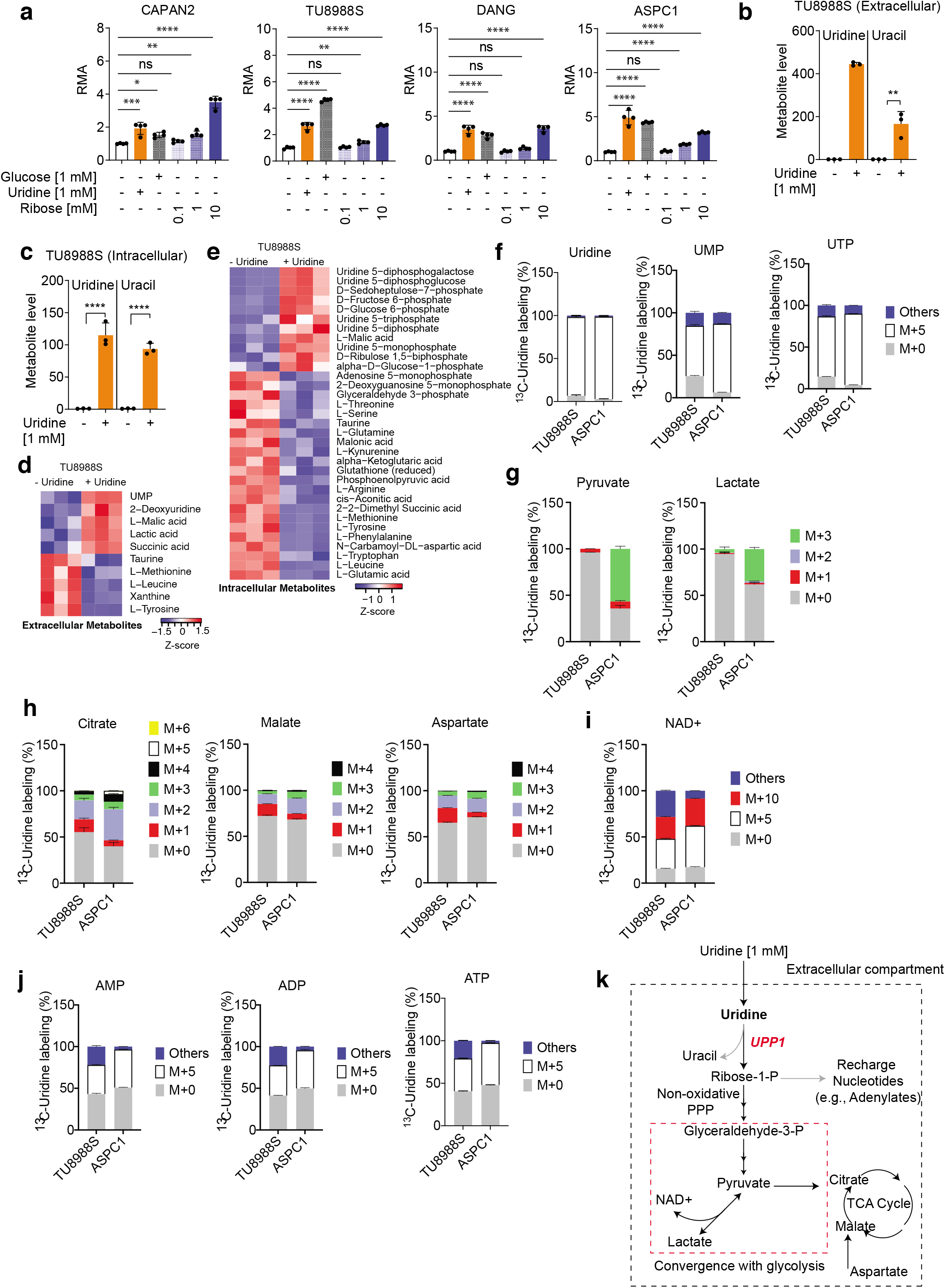
Metabolic profile upon uridine supplementation. **a**. Relative metabolic activity (RMA) of the PDA cell lines supplemented with glucose, uridine or ribose under nutrient-limited culture conditions. Statistical significance was measured by ANOVA. ns = not significant; * = *P*<0.05; ** = *P*<0.01; *** = *P*<0.001, **** = *P*<0.0001. **b-e**. Mass spectrometry metabolomics indicating intracellular metabolite profile after 24 h culture in media supplemented with 1 mM uridine in media with 5% dialyzed FBS. Statistical significance is measured using a Student’s t-test. ns = not significant; * = *P*<0.05; ** = *P*<0.01; *** = *P*<0.001, **** = *P*<0.0001. **f-j**, Mass isotopologue distribution of uridine-derived carbon in the indicated metabolites. M – mass; ‘Others’ – indicate M other than M+0 or M+5, where applicable. **k**, Schematic illustration of uridine utilization based on metabolomics results and known biochemical processes. PPP – pentose phosphate pathway; P – phosphate.

Considering this observation, we employed liquid chromatography mass spectrometry-based metabolomics (Lee et al., 2019) in the UPP1-low cell line PA-TU8988S, and UPP1-high cell line DANG that we used in the initial phases of this work (**Fig. S3a-d**). In PA-TU8988S, supplementation with uridine led to ~200-fold increase in media uracil content consistent with a direct substrate-product conversion (**Fig. 3b**). Further, intracellular uracil almost reached similar levels as uridine (**Fig. 3c**). Uridine supplementation increased lactate secretion (typical aerobic glycolysis/Warburg effect), as well as extracellular malate from the TCA cycle, and uridine derivates UMP and UDP (**Fig. 3d, Fig. S3c**). This elevated level of glycolytic intermediates, uridine derivatives, and TCA cycle intermediates, notably malate, was also observed in the intracellular compartment (**Fig. 3e**) and several more metabolites in these pathways emerged in DANG (**Fig. S3c-d**). Thus, we establish the direct metabolic utility of uridine in the PDA cells. On the other hand, we consistently observed a depleted level of several amino acids, notably methionine, serine, tryptophan and amino acid derivatives kynurenine and glutathione either in the media, intracellular compartment, or both across the two cell lines (**Fig. 3d-e, Fig. S3c-d**). These data could suggest that uridine supplementation promote the utility of amino acids.

To further delineate how uridine is metabolized, we performed mass spectrometry-based metabolomics (Yuan et al., 2019) to analyze the mass isotopologue distribution of isotopically labelled uridine (^13^C_5_-uridine, labeled exclusively on the ribose carbon), using PA-TU8988S, and the UPP1-high uridine high consumer cell line ASPC1, which was more amenable to gene manipulation than DANG. Both cell lines demonstrated high uridine, UMP and UTP labeling, >90%, as indicated by M+5 from the ribose component (**Fig. 3f**). We observed a higher ^13^C labeling in glycolytic intermediates, pyruvate and lactate, in the uridine-high consumer (**Fig. 3g**). Further, there was a considerable labeling of TCA cycle intermediates citrate and malate, and the associated metabolite aspartate (**Fig. 3h**). Uridine-derived ribose-carbon labeled several energy-rich metabolites namely NAD+ (i.e., M+5, M+10), AMP, ADP and ATP (i.e., M+5) (**Fig. 3i-j**), reflecting the direct recycling of the ribose for ribosylation of the adenine base and support redox homeostasis. These data support our model (**Fig. 3k**) illustrating that uridine catabolism converges with glycolysis and fuels both central carbon metabolism and nucleotide biosynthesis in PDA cells.

#### UPP1 directly mediates uridine catabolism in PDA cells to provide ribose

To confirm the regulatory role of UPP1 in uridine catabolism, we knocked out UPP1 using CRISPR-Cas9. We observed that NADH production from uridine catabolism was completely abolished upon UPP1 knockout (UPP1-KO). This was the case in both a UPP1-low (TU8988S, n=4 clonal lines) and a UPP1-high cell line (ASPC1, n=2 clonal lines) (**Fig. 4a**). These data illustrate that cells with lower UPP1 expression are not necessarily less dependent on UPP1. The ability of exogenous uridine to rescue cellular bioenergetics, as read out by ATP, was also suppressed in UPP1 knockout cells grown in nutrient limiting conditions (**Fig. 4b**). These results reveal that the UPP1-mediated catabolism of uridine to uracil and ribose-1-phosphate is indispensable for the utilization of uridine to support reducing potential and proliferation.

**Figure 4.**
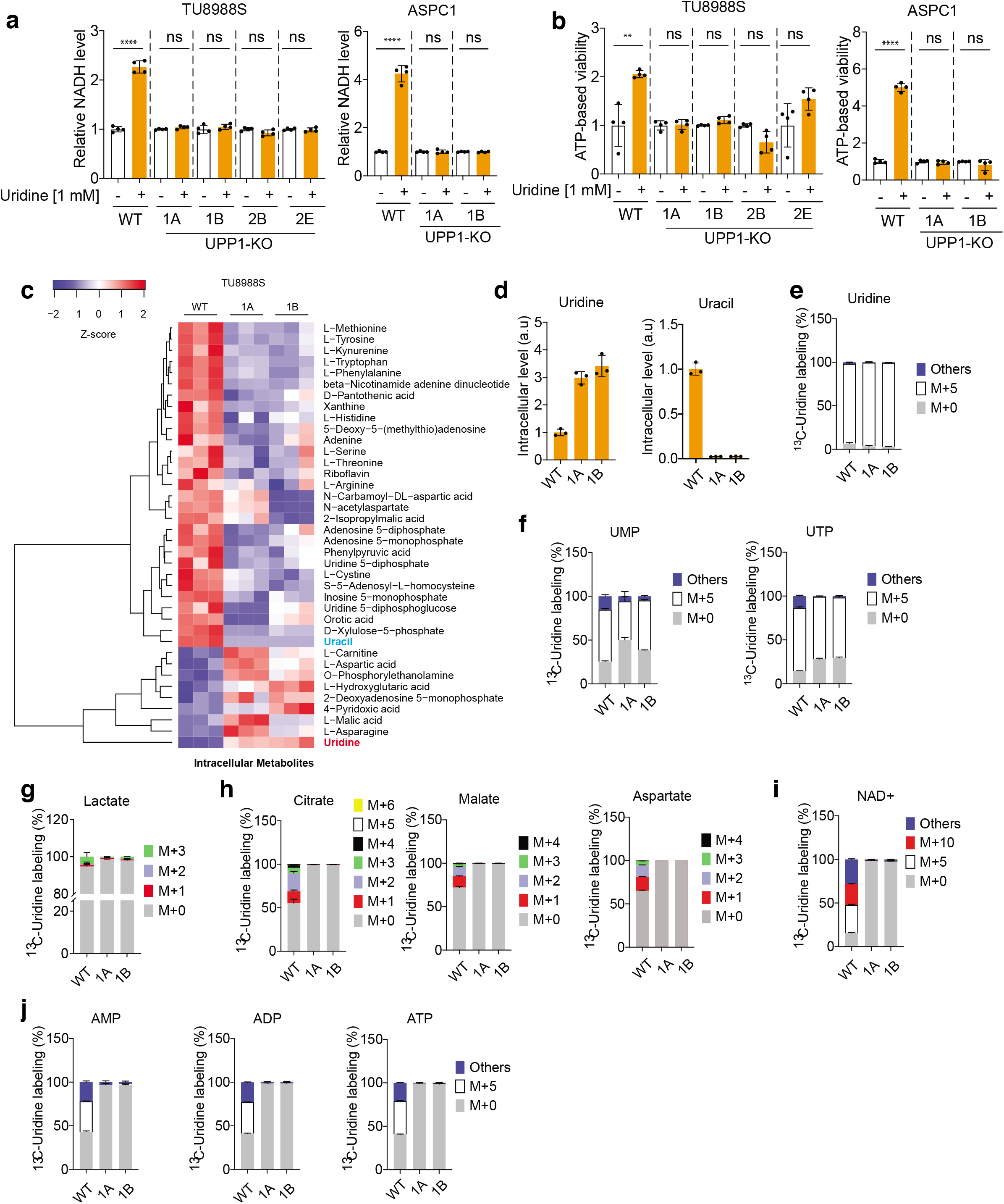
UPP1 mediates uridine catabolism. **a**, Assessment of reducing potential in *UPP1* knockout cell lines versus wild type (WT) supplemented with uridine. A, B, E denotes clones; #1 and 2 denotes sgRNAs; WT – wildtype. Statistical significance is measured using a Student’s t-test. ns = not significant; * = *P*<0.05; ** = *P*<0.01; *** = *P*<0.001, **** = *P*<0.0001. **b**, ATP-based viability assay following uridine supplementation of TU8988S and ASPC1 upon *UPP1* knockout. **c-d**, Mass spectrometry-based metabolomics indicating intracellular metabolite profile after 24 h culture in media supplemented with 1 mM uridine. **e-j**, Metabolomics profiling showing the mass isotopologue distribution of ^13^C_5_-uridine in metabolites involved in glycolysis, the TCA cycle, and nucleotide salvage.

To then determine how UPP1 knockout impacted the metabolism of uridine, we again utilized our metabolomics profiling approach (**Fig. 4c**). Here, we only employed UPP1-KO PA-TU8988S cells because the cell viability of ASPC1 UPP1-KO cells was severely compromised in nutrient-deprived conditions. First, analysis of the metabolomics data from UPP1-KO PA-TU8988S cells revealed several metabolites whose intracellular levels were strongly depleted upon UPP1 knockout (e.g., methionine, tyrosine, kynurenine, serine, arginine, and uridine diphosphate) and also showed an accumulation of malate, aspartate and asparagine (**Fig. 4c**). Importantly, intracellular uridine was high in UPP1-KO cells, confirming that the catabolism of uridine was blocked in these cells. This blockade was further confirmed by the marked depletion of intracellular uracil (**Fig. 4d**), a direct metabolic product of uridine catabolism. Thus, the knockout of UPP1 had a potent impact on uridine-to-uracil conversion as well as various other aspects of central carbon metabolism.

Next, to more precisely determine how UPP1 KO impacted the fate of uridine, we again utilized our isotope tracing metabolomics platform. Ribose-labeled ^13^C_5_-uridine tracing revealed that the intracellular level of labeled uridine was similar in the UPP1-KO and control cell lines (**Fig. 4e**), indicative of uridine uptake at steady-state. Labeling in uridine-derived metabolites, specifically UMP and to a lesser extent UTP, was reduced upon UPP1 knockout (**Fig. 4f**). In addition, flux of uridine (ribose)-derived carbon into glycolysis, as evidenced by labeling of lactate (**Fig. 4g)**, was eliminated in UPP1 knockout cells. Isotope labeling of the TCA cycle-associated metabolites was also depleted (**Fig. 4h**). Furthermore, the recycling of uridine-derived ribose was also completely blocked in the UPP1 knockout cells, as evidenced by the absence of carbon labeling in NAD+ (**Fig. 4i**), and bioenergetic metabolites AMP, ADP, and ATP (**Fig. 4j**). These results provided detailed molecular confirmation that UPP1 directly controls the utilization of uridine in PDA cells.

#### UPP1 drives compensatory uridine metabolism in the absence of glucose to support PDA growth

Our data up to this point illustrate a clear role for the metabolism of uridine-derived ribose through UPP1 to support PDA cell metabolism. Next, we sought to assess if uridine also had an impact on PDA cell proliferation, and the nutrient availability context in which this was relevant. First, we observed that tumors with high UPP1 expression also have a high expression of glycolysis genes, and gene set enrichment analysis showed the upregulation of the glycolytic pathway (**Fig. 5a**). Given our observation that uridine and glucose metabolism converge, we tested whether uridine serves as a nutrient source in the absence of only glucose. In addition, we tested whether glutamine could serve this role, as we have previously shown it to be an important anaplerotic substrate for the TCA cycle (Son et al., 2013). We found that uridine supplementation increased NADH production in glucose and glutamine-free media in three of four PDA cells lines, had a greater impact on NADH production when only glucose was lacking, and had no effect with glucose present (**Fig. 5b**). Importantly, cell proliferation was potentiated by uridine supplementation in glutamine-containing media (**Fig. 5c**). These results are consistent with our biochemical analysis, reflecting that uridine-derived glucose can fuel glycolysis, the TCA cycle, and the nucleotide salvage pathway in place of glucose.

**Figure 5.**
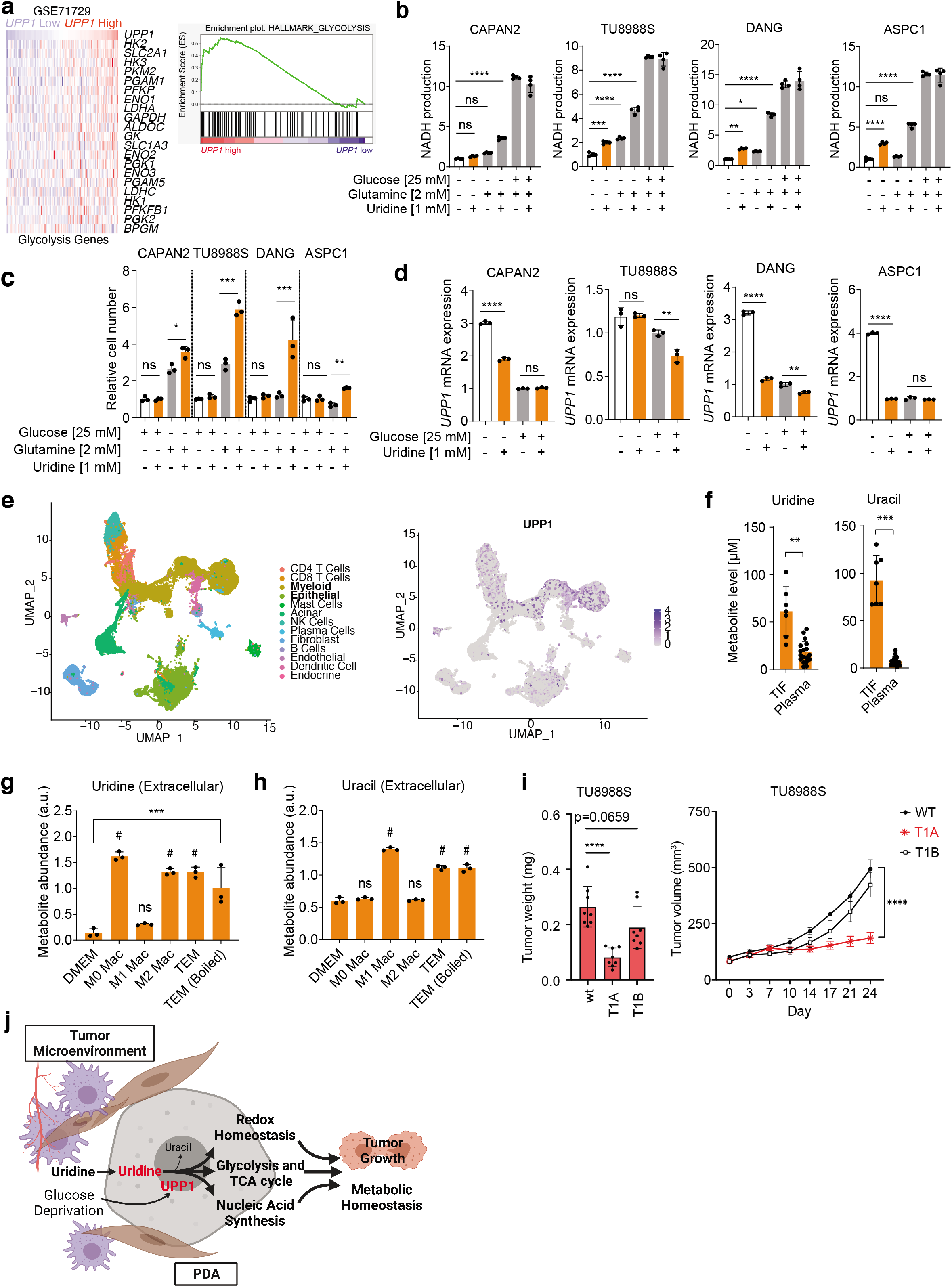
Uridine supports PDA cell growth and metabolic homeostasis during glucose deprivation. **a**, Heatmap showing the expression of glycolysis genes in tumors stratified by *UPP1* expression. On the right, GSEA plots showing the enrichment of glycolysis and hypoxia signatures in UPP1-high tumors. **b**, Biolog assay showing NADH production upon uridine supplementation with or without glucose and glutamine. **c**, CyQUANT proliferation assay upon uridine supplementation with or without glucose and glutamine. **d**, qPCR data showing mRNA expression of *UPP1* in nutrient-deprived media or upon uridine supplementation, after 24 h culture. **e**, Uniform Manifold Approximation and Projection showing *UPP1* expression in tumor epithelial and TME populations, notably myeloid cells in human Single-cell RNA sequencing data. **f**, Metabolomics data showing uridine and uracil levels in tumor interstitial fluid (TIF) relative to the plasma extracted from a previous study (Sullivan et al., 2019). **g-h**, Metabolomics data showing uridine and uracil levels in conditioned media from macrophage subtypes. TEM – tumor educated macrophages. **i**, Tumor volume of *UPP1* KO in the UPP1 high cell line (WT n = 7; UPP1 KO n = 8 tumors per clone) ns = not significant; * = *P*<0.05; ** = *P*<0.01; *** = *P*<0.001, **** = *P*<0.0001. **j**, Graphical illustration of the connection between tumor microenvironment, uridine and UPP1-catalized metabolism, and PDA growth.

Because of the impact of glucose availability on the utility of uridine-derived ribose, we hypothesized that glucose availability could signal a transcriptional change to regulate UPP1 expression. Indeed, the removal of glucose from the culture media induced a strong increase in UPP1 expression (**Fig. 5d**). However, the increase in UPP1 expression was reversed in media lacking glucose but containing 1mM uridine. Thus, both exogenous uridine and glucose exert control over UPP1 expression. These results support a model in which nutrient deprivation leads to transcriptional upregulation of UPP1.

Considering the utility of uridine as a novel nutrient input in PDA cell metabolism and growth, we wondered if uridine is abundant in the tumor microenvironment and what its sources might be. First, we assessed the expression of UPP1 in our published single-cell RNA sequencing data (Steele et al., 2021). UPP1 expression was high in both the tumor epithelial and myeloid cells, which consist mainly of macrophages (**Fig. 5e**). Using a previously published dataset of metabolites in tumor interstitial fluid (TIF) (Sullivan et al., 2019), we also found that uridine and uracil levels were enriched in TIF, relative to serum (**Fig. 5f**), reaching micromolar levels and suggesting greater availability in the tumor microenvironment. Indeed, we also observed that macrophages that were differentiated *in vitro* and polarized to a tumor-educated fate with PDA cell conditioned media release uridine and uracil (**Fig. 5g-h**), indicating a potential source in the pancreatic TME. Lastly, the knockout of UPP1 impaired tumor growth in murine subcutaneous xenograft model of PDA (**Fig. 5i**). Taken together, our data suggest that UPP1 is a potential mediator of compensatory metabolism and growth in nutrient-limited conditions and that macrophages are capable of secreting uridine into the extracellular compartment (**Fig. 5j**).

## Discussion

It is well acknowledged that understanding the metabolic needs of PDA can help to determine the drivers of the disease aggressiveness and therapeutic resistance, and unravel new opportunities for therapy (Biancur and Kimmelman, 2018; Halbrook and Lyssiotis, 2017). Despite this, there is a limited knowledge of the diversity of the nutrients utilized by PDA cells. To address this shortcoming, we employed a high throughput assay of nutrient utilization in a panel of diverse, well characterized PDA cell lines. We found that PDA cells use extracellular uridine-derived ribose through UPP1-mediated catabolism in nutrient-limited conditions to support cellular metabolism and growth. Our metabolomics profiling, which includes stable isotope tracing of uridine-ribose carbon, resolved the metabolic fate of uridine – namely, a contribution to glycolysis, TCA cycle, NAD+, and energy-rich products including ATP. Further, we showed that UPP1 is itself controlled by nutrient availability in PDA cells, consistent with its role in supporting compensatory metabolism. Finally, we demonstrate that uridine is enriched in the pancreatic TME, where it may serve as a novel nutrient source, and we illustrate the potential relevance of UPP1 using patient transcriptomics data, single cell RNA sequencing datasets and tumor growth studies.

Nutrients in the TME used by PDA cells can be derived from a variety of sources, including other cell types. We previously demonstrated the direct exchange of alanine from pancreatic cancer associated fibroblasts to PDA cells (Sousa et al., 2016) by profiling metabolites depleted from supporting-cell conditioned media using a metabolomics-based approach. While high-throughput, this method simplifies the complex metabolic interactions of the TME to a two-cell type system. Others have used the Biolog Phenotype Microarrays, showing metabolic shifts corresponding to the epithelial-mesenchymal transition in breast cancer stem-like cells, as well as following melittin treatment in cisplatin-sensitive and -resistant ovarian cancer cells (Alonezi et al., 2016; Cuyàs et al., 2014). We uniquely utilize this system to first obtain source-agnostic information about the nutrients utilized by PDA cells, before conducting a targeted analysis of potential cell types that may provide exogenous uridine.

Uridine and uracil are both enriched in the TIF of murine pancreatic tumors, relative to serum. Furthermore, our *in vitro* analysis of the extracellular metabolome revealed micromolar release of uridine from macrophages exhibiting either an anti-inflammatory phenotype or those polarized by the conditioned media of PDA cells. Macrophages are the most abundant immune cell type in PDA tumors and can constitute upwards of 40% of the tumor by content (Halbrook et al., 2019). Thus, we put forth that the increased uridine in the pancreatic TME may be derived from tumor associated macrophages. Further, additional recent precedent for nucleic acid (e.g. inosine) consumption in melanomas by both cancer and CD8^+^ effector T cells leads us to the conclusion that nucleoside metabolism in solid tumors is complex but therapeutically tractable (Wang et al., 2020). There are indeed a diverse range of cell types that can utilize uridine for various purposes (Choi et al., 2006, 2008; Geiger and Yamasaki, 1956), and this is a provocative area for future study.

Our finding that exogenous uridine supplements cellular energetic and biosynthetic needs in the absence of sufficient glucose aligns with other relevant studies. Though not previously shown to be a cancer-supporting mechanism, uridine also rescues cellular nutrient stress due to glucose deprivation in astrocytes and neurons (Choi et al., 2006, 2008; Geiger and Yamasaki, 1956). In cancer cells, and potentially neurons and astrocytes, the ribose ring fuels both energetic and redox metabolism via routing through the PPP and subsequent oxidative catabolism or, alternately, anabolism into biomass for cellular proliferation, e.g., via the nucleic acid salvage pathway. We and others have shown that this mechanism is UPP1-dependent, the expression of which is dependent on nutrient availability (Choi et al., 2008). Both glucose and uridine-derived ribose participate in the support of the energy charge of the cells by facilitating ATP generation, which ratiometrically with AMP controls the activation of AMPK signaling (Garcia and Shaw, 2017; Xiao et al., 2011). p53 interacts with AMPK signaling and is a known transcriptional repressor of UPP1 (Maddocks and Vousden, 2011; Zhang et al., 2001). Therefore, in line with our observation of high UPP1 in glucose-free media, we suggest that glucose deprivation and lowered ATP trigger likely adaptive metabolism in PDA.

The upregulation of nucleoside usage in diverse cell types under nutrient deprivation (Jurkowitz et al., 1998; Litsky et al., 1999; Wang et al., 2020), especially immune and PDA cells, implies metabolic competition that may contribute to immunosuppression and tumor progression. UPP1 therefore represents a double-edged sword with respect to PDA treatment. Namely, UPP1 activity is required for conversion of fluoropyrimidine prodrugs into active anti-cancer therapies, particularly the 5-fluorouracil prodrug capecitabine, a drug in the multi-drug combination and front-line therapy FOLFIRINOX used to treat pancreatic cancer (Cao et al., 2002; Longley et al., 2003). Future studies may therefore endeavor to determine the significance of proteins downstream of UPP1 and other nucleoside phosphorylases that prevent ribose1-phosphate generation, while not hindering 5-fluorouracil-related drugs.

## Materials and Methods

### Cell Culture

20 human PDA cell lines and 2 from normal human pancreas tissue were purchased from the American Type Culture Collection (ATCC). hPSC cell lines were a kind gift from Rosa Hwang (MD Anderson Cancer Cencer, Texas, USA). The confirmation of the identity of cell lines was established with STR profiling, and lines were routinely tested for mycoplasma using MycoAlert (Lonza, LT07-318). All cell lines were cultured in high-glucose DMEM (Gibco, 11965092) supplemented with 10% fetal bovine serum (FBS, Corning, 35-010-CV) at 37°C and 5% CO_2_. Phosphate buffered saline (PBS, Gibco, 10010023) was used for cell washing steps unless otherwise indicated.

### Biolog Metabolic Assay

In the initial phenotypic screen, the 22 cell lines were grown in proprietary 96-well PM-M1 and PM-M2 plates (Biolog, 13101 and 13102). The relative metabolic activity (RMA) from substrate catabolism in the cells was measured using Biolog Redox Dye Mix MB. Briefly, the cell lines were counted, and their viability assessed using Trypan Blue Dye (Invitrogen, T10282). The cells were then washed 2x with Biolog Inoculating fluid IF-M1 (Biolog, 72301) to remove residual culture media. Then, a cell suspension containing 20,000 cells per 50uL was prepared in Biolog IF-M1 containing 0.3mM glutamine and 5% dialyzed FBS (dFBS) (Hyclone GE Life Sciences, SH30079.01) and plated into PM-M1 and PM-M2 96-well plates at 50 μL and 20,000 cells per well. Plates were incubated for 24 hours at 37°C and 5% CO2, after which time 10 μL Biolog Redox Dye Mix MB (Biolog, 74352) was added to each well. Plates were sealed to prevent the leakage of CO_2_. The reduction of the dye over time was measured as absorbance (A590-A750) using the OmniLog® PM-M instrument (Biolog, 93171) for 74.5 hours at 15 minutes intervals. The generated data were analyzed using *opm* package in R statistical programming tool (version 3.5.2). After quality checks of the microarray results, the maximum metabolic activity per cell line was taken as its main readout for substrate avidity. Heatmap visualization of the data was plotted using *heatmap2* package in R.

### CyQUANT Proliferation Assay

The PDA cells were seeded at 2,000 cells per well in a 96-well plate (Corning, 3603) and in regular DMEM. The next day, the culture media was removed, followed by a gentle 1x wash with PBS. Treatment media was then applied, and the cells incubated at 37°C and 5% CO_2_ until they reached ~70% confluence. Media was then carefully aspirated, and the plate with cells frozen at −80°C for at least 24 hours to lyse all cells. To prepare the lysis buffer and DNA dye, CyQUANT Cell Lysis Buffer and CyQUANT GR dye (Invitrogen, C7026) were diluted in water at 1:20 and 1:400, respectively. The frozen cells were then thawed and 100 μL of the lysis buffer was added to each well. Thereafter the plate was covered to prevent light from inactivating the GR dye and was placed on an orbital shaker for 5 minutes before measurement. Fluorescence from each well, indicating GR dye binding to DNA, was then measured utilizing a SpectraMax M3 Microplate reader at an excitation wavelength of 480nm and an emission wavelength of 520nm.

### ATP-based viability assay

The cell lines were seeded in quadruplicates at a density of 2,500 – 5000 cells in 100 μL DMEM per well of white-walled 96-well plates (Corning/Costar, 3917). Next day, the media was aspirated, each well was washed with 200 μL PBS after which treatment media was introduced. At the end of the experiment duration (i.e., 72 h or as otherwise indicated), cell viability was determined with CellTiter-Glo 2.0 Cell Viability Assay Kit (Promega, G9243) and the luminescence quantified using a SpectraMax M3 Microplate Reader.

### UPP1 CRISPR/Cas9 knockout

The expression vector pspCas9(BB)-2A-Puro (PX459) used for UPP1 CRISPR/Cas9 construct was obtained from Addgene (Plasmid #48139). The plasmid was cut using the restriction enzyme *BbsI* followed by the insertion of UPP1 sgRNAs (**Table 2)** as previously described (Ran et al., 2013). For transfection, PDA cells were seeded at 150,000 cells per well in a 6 well plate one day prior. The cells were transfected with the plasmid pSpCas9-UPP1 using Lipofectamine 3000 Reagent (Invitrogen, L3000001) in the following proportion: 1μg of DNA, 2μL Lipofectamine, and 2 μL P3000 reagent per 1mL of transfection media. After 24 h, the selection of successfully transfected cells was commenced by culturing the cells with 0.3 mg/mL puromycin in regular DMEM. The puromycin-containing media was replaced every two days until selection was complete as indicated by the detachment of all non-transfected cells. Thereafter, the successfully transfected cell lines were expanded and clonally selected after serial dilution.

**Table 2.**
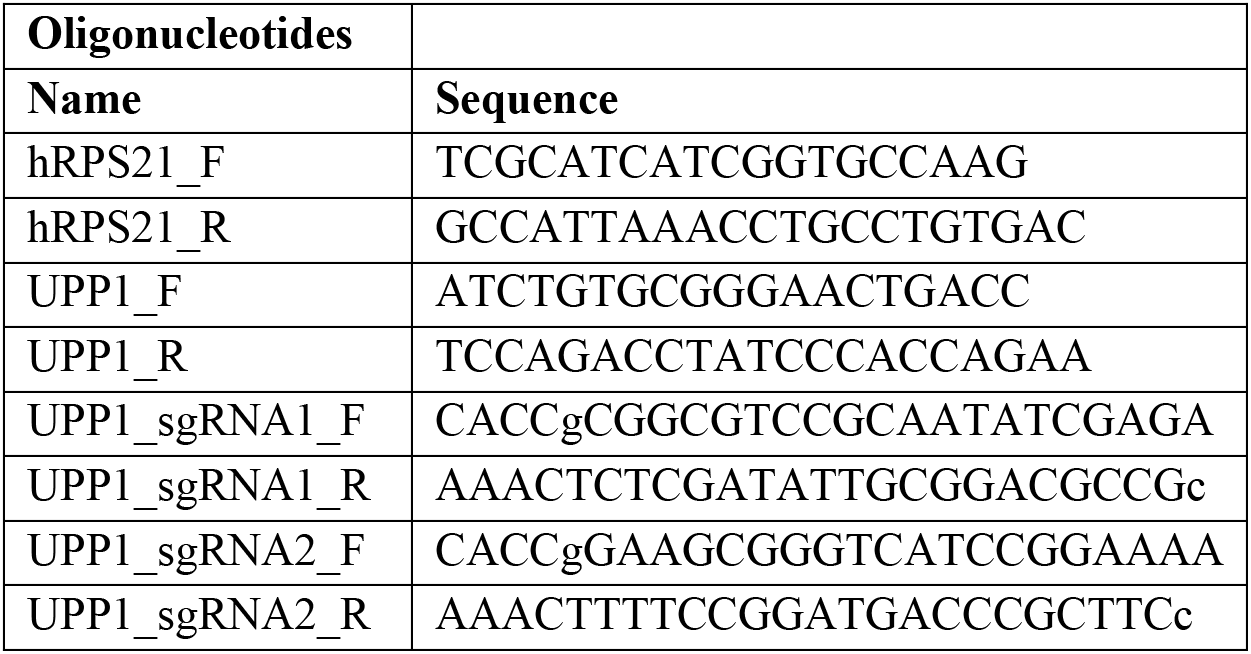
Oligonucleotides.

### Quantitative Reverse-Transcriptase Polymerase Chain Reaction

RNA samples were isolated using the RNEasy Plus Mini Kit (QIAGEN, 74134) according to the manufacturer’s instructions. RNA purity was assessed using a NanoDrop One (ThermoFisher Scientific, ND-ONE-W). The RNA samples (1μg) were thereafter reverse transcribed to cDNA using the iScript cDNA Synthesis Kit (Bio-Rad, 1708890) according to the accompanying instructions. qPCR was then performed on the QuantStudio 3 Real-Time PCR System (ThermoFisher Scientific, A28136) using Power SYBR Green PCR Master Mix (ThermoFisher Scientific, 4367659). Primer sequences are in **Table 2**. The gene expression was calculated as delta CT and *RPS21* was used as a housekeeping gene.

### Metabolomics and stable isotope tracing

For extracellular and unlabeled intracellular metabolomics profiling, the PDA cells were seeded in a 6-well plate at 400,000 cells per well in DMEM culture media. A parallel plate for protein estimation and sample normalization was also set up with the same number of cells. After overnight incubation, the culture media was aspirated off and replaced with ‘treatment media’ with or without 1 mM uridine. The cells were then cultured for a further 24 hours. Thereafter, for extracellular metabolites, 200 μL of media was collected from each well into a 1.5 mL Eppendorf tube and to that 800 μL ice-cold 100 % methanol was added. For intracellular, the remaining media was aspirated off and the samples washed 1x with 1mL cold PBS before incubation with 1 mL ice cold 80% methanol on dry ice for 10 minutes. Thereafter, cell lysates were collected from each well and transferred into separate 1.5 mL Eppendorf tubes. The samples were then centrifuged at 12,000 *g* and the volume of supernatant to collect for each experimental condition was then determined based on the protein concentration of the parallel plate. The collected supernatants were dried using SpeedVac Concentrator, reconstituted with 50% v/v methanol in water, and analyzed by mass spectrometry. For stable isotope tracing, we used ^13^C_5_ (ribose) labeled uridine (Cambridge Isotope Laboratories, CLM-3680-PK) supplemented at 1mM. Briefly, cells were cultured overnight in regular media. Next day, cells were washed 1x followed by the introduction of glucose-free media supplemented with labelled uridine. In parallel, a similar experiment was set up for unlabeled uridine. The cell lines were then cultured in the uridine-supplemented media for 24 h, followed by sample collection as with unlabeled intracellular metabolomics above. Samples for unlabeled and ^13^C-(ribose) uridine tracing analyses were run on targeted liquid chromatography tandem mass spectrometry (LC-MS/MS) and processed as previously described (Nelson et al., 2020).

### Mouse xenograft tumors

Animal studies were performed in accordance with the guidelines of Institutional Animal Care and Use Committee (IACUC) and approved protocol. Female athymic nude mice NU/J (Stock No: 002019, The Jackson Laboratory) of ~6 weeks old were maintained in the facilities of the Unit for Laboratory Animal Medicine (ULAM) under specific pathogen-free conditions. On the day of tumor cell injection, PA-TU-8988S wildtype (WT) and UPP1 knockout clonal cells were harvested from culture plate according to normal cell culture procedures. The cells were counted, washed 1x with PBS and resuspended in 1:1 solution of serum free DMEM (Gibco, 11965-092) and Matrigel (Corning, 354234). Mice were subcutaneously injected on both flanks with 500,000 cell lines in a 100 μL final volume. Once tumors became palpable, tumor sizes were monitored twice/week using a digital caliper until end point. Tumor volume was calculated as V = 1/2(length × width^2^).

### PDA dataset analysis

The human PDA microarray datasets with accession numbers GSE71729 (n=46 normal pancreas vs 145 tumor tissues) and GSE62452 (n=61 non-tumoral vs 69 tumor tissues) were obtained from NCBI GEO (Barrett et al., 2013). Differential gene expression between PDA and non-tumors were performed in R. Kaplan Meier (KM) overall survival (log-ranked test) was performed after splitting the tumor samples per dataset into UPP1 high and low subsets. For KM, the following tumor datasets were used: GSE71729 (n=145), the cancer genome atlas (TCGA) data (n=146), international cancer genome consortium (ICGC, n=267), and Puleo et al. (n=288). The iKras mice data was obtained from NCBI GEO under the accession number GSE32277.

### CCLE gene analysis and PDA tumor data stratification

Following the determination of uridine higher and lower consumer PDA cell lines from the Biolog assay, the gene expression data of these two subsets were extracted from the cancer cell line encyclopaedia (CCLE, GSE36133). The subsets were then compared using *limma* package in R to determine the differentially expressed genes in uridine-high consumers relative to lower consumers. For the tumor stratification, samples in the dataset GSE71729 (n=145) were ranked into UPP1-high and low groups and compared as above to determine the genes differentially expressed in UPP1 high tumors.

### Pathway analyses

Pathway analysis were performed using DAVID functional annotation platform (https://david.ncifcrf.gov/, v 6.8), or the gene set enrichment analysis (GSEA, v 4.0.3) with GSEAPreranked option. Ranking of the genes was based on the product of the logFC and −log(p-value). GSEA was run with default parameters, except gene set size filter set at min=10. Gene ontology analyses were performed with DAVID.

### Statistics

Statistics were performed either with GraphPad Prism 8 (GraphPad Software Inc.) or using R. Two group comparisons were made using the two-tailed Student’s t-test. The error bars in all graphs represent the mean±standard deviation, and significance annotations are noted in each figure. *P* < 0.05 was considered statistically significant unless indicated otherwise. Heatmaps were plotted with Heatmap2 package in R.

## Acknowledgments

The authors would like to thank Dr. Barry Bochner (Biolog, Inc., USA) for providing the OmniLog instrument and helping with the data analysis. We also thank Serena Chan for support with Biolog data collection and analysis. Further, we thank members of the Sadanandam and Lyssiotis labs and the entire Pancreatic Disease Initiative at the Rogel Cancer Center, University of Michigan, for their insightful comments and discussions.

## Supplementary Figure Legends

**Figure S1. The Biolog assay and the measured parameters**

**a**, Schematic overview of representative data from the Biolog Phenotype Microarray, as measured by the Omnilog Machine. Parameters were derived from the spline-fit curve.

**b-c**, Boxplot distribution of estimations of the AUC parameters in all cell lines, with each point representing a nutrient source in the PM-M1 plate (b) and PM-M2 plate (c).

**d**, Correlation between the AUC and A parameter estimations for every nutrient source and cell line combination in the PM-M1 plate.

**e**, Heatmap showing the stratification of the cell lines based on consumption of metabolites in PM-M2 plate.

**f**, Heatmap showing metabolites differentially consumed by the PDA cell lines analyzed using Biolog assay PM-M2 plates.

**g**, Heatmap showing the topmost PM-M1 plate metabolites consumed by most of the tested cell lines.

**Figure S2. *UPP1* expression correlates with uridine**

**a**, Correlation of *UPP1* expression with measured uridine consumption (left). On the right, qPCR validation of UPP1 expression in the cell lines identified as uridine low and high consumers from the Biolog assay.

**b,** Correlation of *UPP1* expression with other nucleosides.

**c**, Gene ontology ‘cellular component’ altered in *UPP1*-high cell lines.

**d**, *Upp1* expression in iKras cell lines in the dataset GSE32277.

**e**, GSEA plots depicting pathways upregulated in UPP1 high tumors from GSE71729 dataset.

**f**. Heatmap showing tricarboxylic acid (TCA) cycle genes differentially expressed in tumors stratified by *UPP1* expression. Correlation analysis were Spearman’s.

**Figure S3. Uridine supports PDA metabolism**

**a-d**, Intracellular and extracellular metabolite profile of PDA cell line DANG cultured in media with or without 1 mM uridine for 24 h. Statistical significance was assessed by t-test. ** = *P*<0.01; *** = *P*<0.001.

