## Supplementary Figures for "Nutrient profiling reveals extracellular uridine as a fuel for pancreatic cancer through uridine phosphorylase 1"

### **Figure S1.** The Biolog assay and the measured parameters

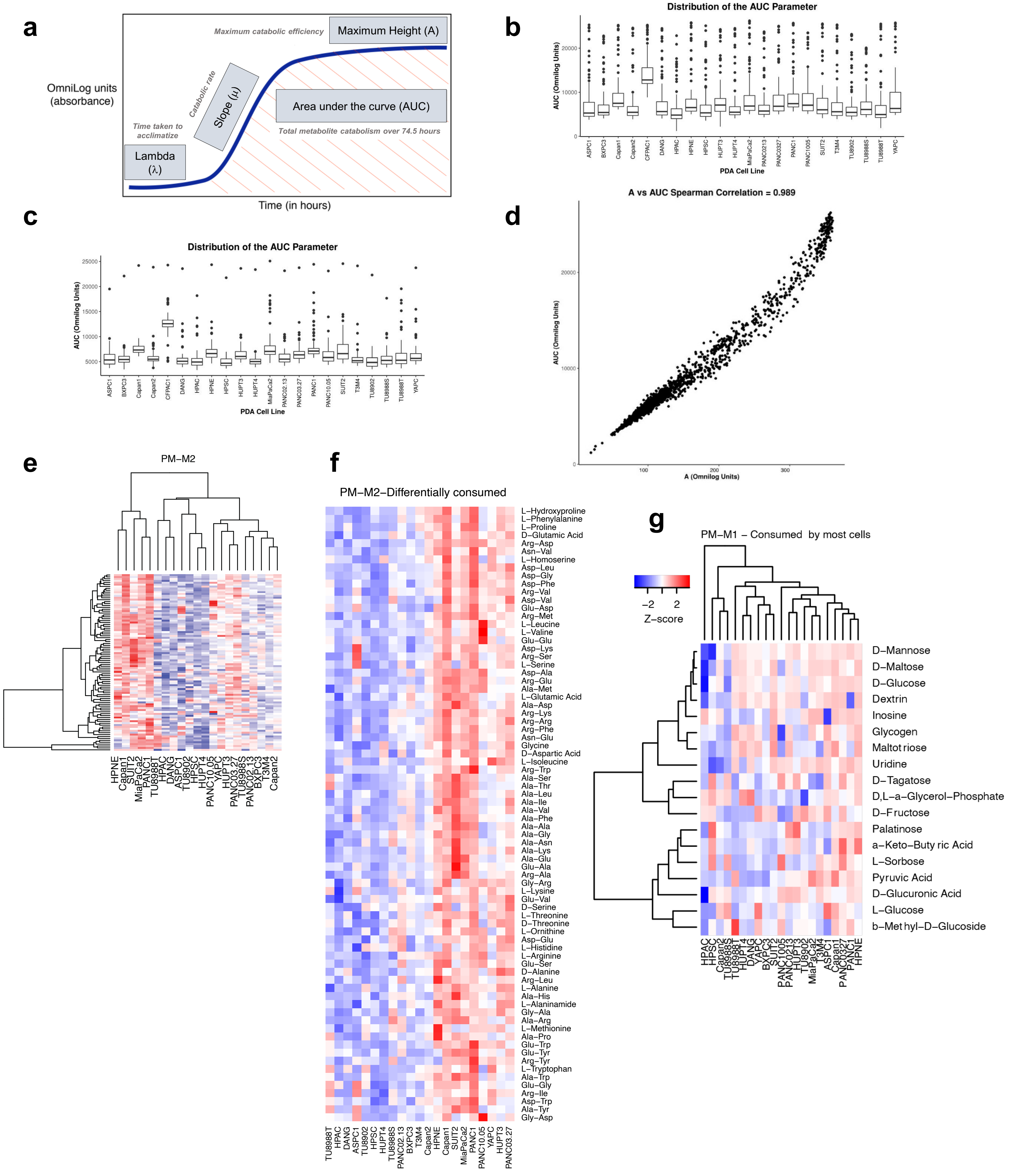

**Figure S2.** UPP1 expression correlates with uridine

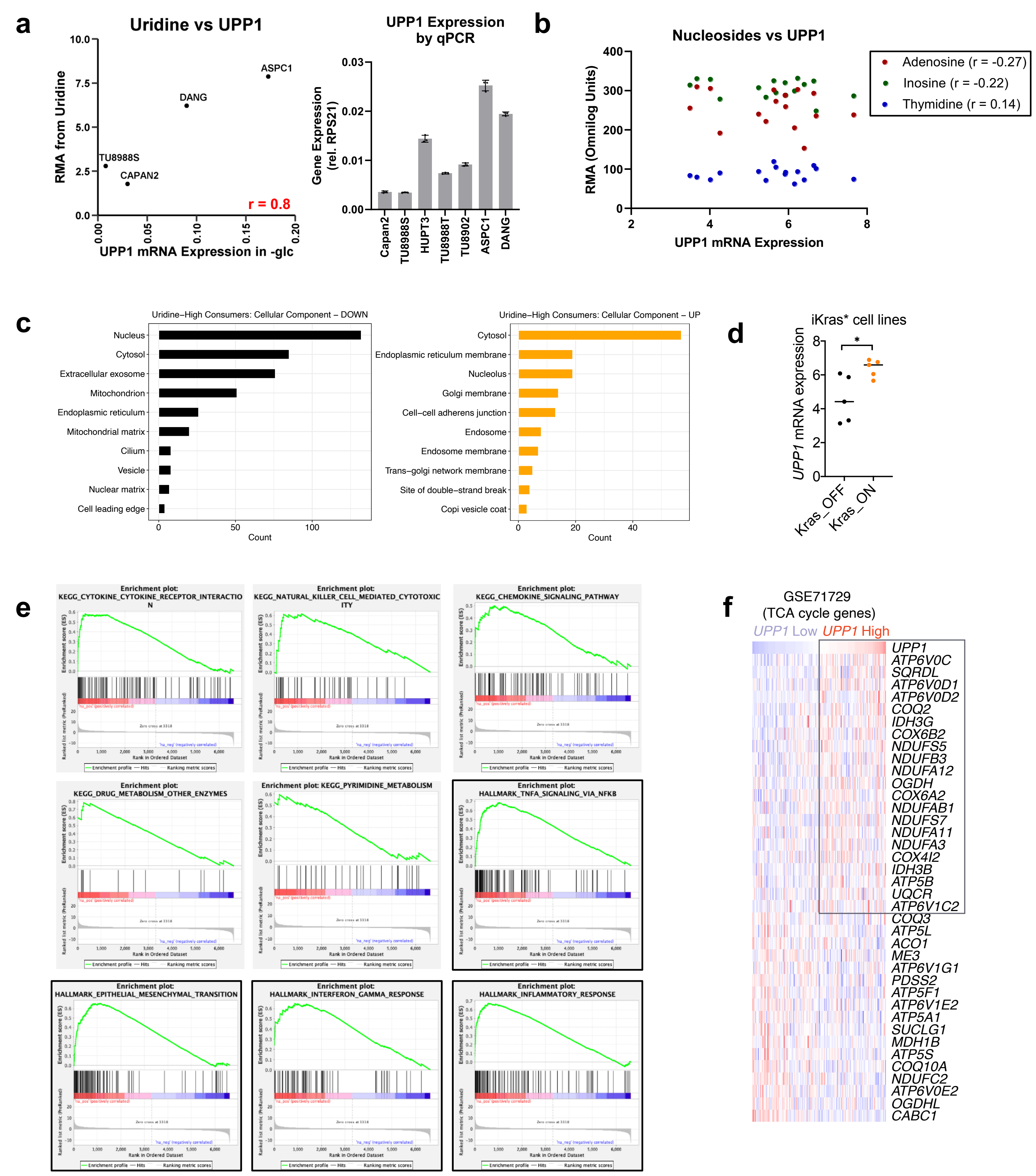

### **Figure S3.** Uridine supports PDA metabolism

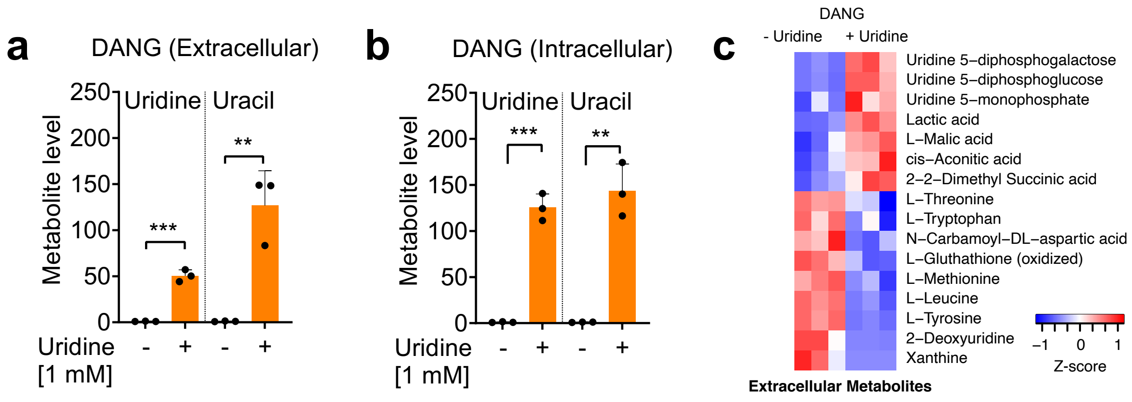


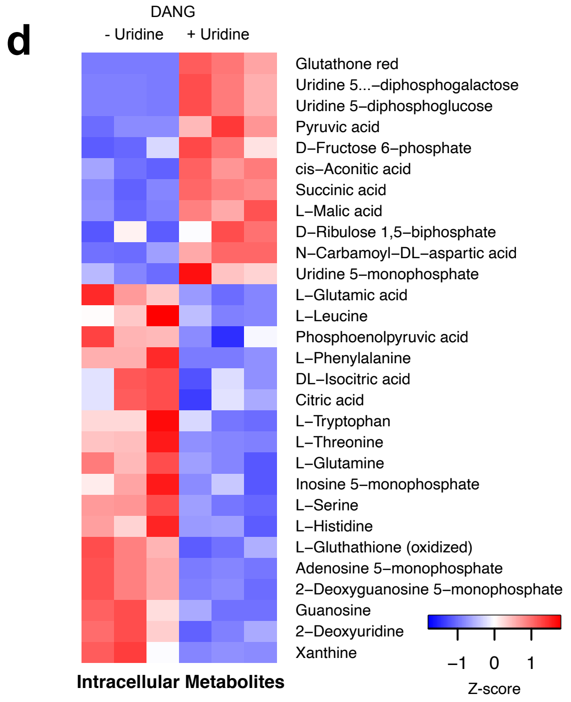

**Supplementary Table 2.** Significant high confidence metabolite-gene correlations

| *r* | Metabolite | Gene.Symbol | Gene.Name |
|---|---|---|---|
| 0.947 | Succinamic Acid | KCNC1 | potassium voltage gated channel, Shaw-related subfamily, member 1 |
| 0.875 | Turanose | B3GALT2 | UDP-Gal:betaGlcNAc beta 1,3-galactosyltransferase, polypeptide 2 |
| 0.868 | D-L-Lactic Acid | THEM4 | thioesterase superfamily member 4 |
| 0.844 | Succinamic Acid | CHST10 | carbohydrate sulfotransferase 10 |
| 0.838 | Uridine | PECI | peroxisomal D3,D2-enoyl-CoA isomerase |
| 0.835 | Mono-Methyl Succinate | KCNC1 | potassium voltage gated channel, Shaw-related subfamily, member 1 |
| 0.826 | Butyric Acid | SLC1A7 | solute carrier family 1 (glutamate transporter), member 7 |
| 0.824 | D-Galactose | SAT2 | spermidine/spermine N1-acetyltransferase family member 2 |
| 0.821 | a-D-Glucose-1-Phosphate | CBR1 | carbonyl reductase 1 |
| 0.821 | Turanose | SARDH | sarcosine dehydrogenase |
| 0.818 | D-Fructose-6-Phosphate | SLC25A39 | solute carrier family 25, member 39 |
| 0.818 | Uridine | UPP1 | uridine phosphorylase 1 |
| 0.815 | Butyric Acid | SLC25A45 | solute carrier family 25, member 45 |
| 0.809 | Butyric Acid | ACBD5 | acyl-Coenzyme A binding domain containing 5 |
| 0.808 | Pyruvic Acid | ABCG4 | ATP-binding cassette, sub-family G (WHITE), member 4 |
| 0.806 | Acetoacetic Acid | FAAH | fatty acid amide hydrolase |
| 0.805 | Turanose | PIK3R6 | phosphoinositide-3-kinase, regulatory subunit 6 |
| -0.803 | D-L-a-Glycerol-Phosphate | IDH3A | isocitrate dehydrogenase 3 (NAD+) alpha |
| -0.809 | D-Glucose-6-Phosphate | DAK | dihydroxyacetone kinase 2 homolog (S. cerevisiae) |
| -0.814 | Glycogen | ASPG | asparaginase homolog (S. cerevisiae) |
| -0.815 | Uridine | ETFA | electron-transfer-flavoprotein, alpha polypeptide (glutaric aciduria II) |
| -0.818 | a-Keto-Glutaric Acid | GNS | glucosamine (N-acetyl)-6-sulfatase (Sanfilippo disease IIID) |
| -0.828 | D-Fructose | KCNK10 | potassium channel, subfamily K, member 10 |
| -0.838 | D-L-a-Glycerol-Phosphate | CHST14 | carbohydrate (N-acetylgalactosamine 4-0) sulfotransferase 14 |
| -0.856 | D-Glucuronic Acid | ATP5SL | ATP5S-like |
| -0.862 | D-Fructose | MAN2A2 | mannosidase, alpha, class 2A, member 2 |
